# Effects of *Tilia tomentosa* on sleep architecture and circadian rhythm

**DOI:** 10.1101/2025.08.08.669271

**Authors:** Antonio Di Soccio, Martina Canova, Nicola Rizzo, Claudia Lodovichi

## Abstract

*Tilia* extract has been used for centuries as a sedative and hypnotic compound. The impact of *Tilia* on sleep architecture and circadian rhythm remained, however, elusive. Here, we addressed these open questions by analysing the behaviour and the EEG signals recorded in freely moving mice fed an enriched diet with *Tilia* extracts. We found that *Tilia* significantly increased the amount of sleep, mostly during the dark hours, which correspond to the most active phase in nocturnal animals, such as mice. Furthermore, in darkness, the length of wake episodes did not increase as in controls. Spectral analyses of the EEG showed that the features of the sleep and wake signals remained unaltered upon *Tilia* treatment. Interestingly, the power in the gamma band resulted significantly decrease while the theta band power resulted increased in treated mice. Altogether, our data demonstrated a clear effect of *Tilia* on sleep architecture, highlighting its dependency on the circadian rhythm.

## Introduction

Anxiety and sleep disorders, which are among the most frequent symptoms in psychiatric and neurological disorders, affect a large portion of the population. The World Health Organization (WHO) estimates that 970 million individuals around the world suffer from these disorders (WHO 2022). Sleep is essential for the overall well-being and health. Sleeplessness is accompanied by altered cognitive functions, increased irritability and anxiety, along with negative emotional states that compromise decision-making ability, reduce work productivity and increase the risk of incidents of various types^1,2^. The heavy burden of symptoms associated with the lack of sleep, prompts a large and increasing use of drugs that favor relaxation and sleep, despite the harmful side-effects they could exert, especially with prolonged use. For this reason, since 2002 WHO has developed and launched the traditional medicine strategy to support and implement the utilization of plants to strengthen the use of traditional medicine to help maintain and promote human health.

Among the plants employed as remedies for human maladies, *Tilia* extracts have been used for centuries in traditional medicine as sedative and tranquilizer compounds. The efficacy of these compounds prompted several studies on mice, aiming at dissecting their mechanism of action.

Behavioral studies showed that mice treated with *Tilia* maintained normal locomotor activity but exhibited a significant increase in immobility time, reduced rearing and other behavioral features that suggest a clear sedative effect^3–5^. In addition, some extracts and several isolated flavonoid fractions from different species of *Tilia* produced an anxiolytic effect on mice tested with the elevated plus maze. In this task, the mice spent more time in and made more frequent crossings of the open arms – an area that typically induces fear or discomfort in mice under normal circumstances^6^. These findings suggest that the *Tilia*-treated mice showed lower anxiety levels, which manifested as increased exploratory behavior. In addition, *Tilia* extracts induced an increase in head diggings in the hole board test and exploration time in the open field test, indicating a reduced level of uneasiness in treated animals that were more inclined to scouting around compared to control, non-treated mice^7^.

It is reasonable to assume that the anxiolytic activity of *Tilia* is due to an effect on GABAergic transmission. GABA, the major inhibitory neurotransmitter in the mature brain, operates via GABA_A_ receptors to trigger fast inhibition, which has a key role in modulating the inhibitory tone, the excitation-inhibition balance and the synchrony among neurons^8^. GABA is also essential in regulating the sleep-wake cycle and sleep architecture^9^.

The molecular mechanism underlying the anxiolytic action of *Tilia* was demonstrated by electrophysiological recordings^10^. In this study, Allio et al. showed that acute application of *Tilia* extracts to hippocampal neurons in culture activated a chloride flux into hippocampal neurons similar to the one elicited by GABA_A_ receptors activation. Most of this effect (about 90%) was blocked by bicuculline and picrotoxin (antagonists of the GABA_A_ receptors), while the remaining 10% was prevented by the benzodiazepine antagonist flumazenil, indicating that *Tilia* extracts act via GABA_A_ receptor activation. The inhibitory action of Tilia via GABA_A_ receptor activation was further demonstrated by the fact that *Tilia* reduced spontaneous neuronal activity and disrupted the synchrony of cultured hippocampal neurons^10^. In this scenario, it is worth noticing that numerous evidence indicates that flavonoids present in several plants, including Tilia, act as ligands of GABA_A_ receptors.^4, 8, 11.^

Despite the well-documented sedative and hypnotic effects attributed to *Tilia*, the impact of *Tilia* extracts on EEG signal features of the sleep-wake cycle and sleep architecture, remains elusive. Here, we addressed this open question performing chronic EEG recordings in freely moving mice treated with extracts from *Tilia Tomentosa* (from now on indicated as *Tilia*). We found that *Tilia* significantly increased the sleeping time and induced a concurrent reduction in wake. Noteworthy, these effects were restricted to the dark phase, which corresponds to the active phase of mice, suggesting a circadian dependency of *Tilia*’s action. In addition, the length of the periods of wake (indicated as bouts of wake) during darkness was shorter in treated subjects than in controls. Consistent with a sedative and hypnotic effect of *Tilia* extracts, treated mice exhibited a reduced number of fast awakenings with respect to controls. To examine the impact of *Tilia* treatment on neuronal oscillatory activity, we conducted EEG spectral analysis. We found that during wakefulness, *Tilia* treatment was associated with a reduction in gamma-band power, while in REM sleep, spectral power increased around 8 Hz, consistent with enhanced theta activity. Again, these effects were confined to the dark (active) phase.

Altogether, our data show that the sedative and hypnotic effects of *Tilia* are supported by distinct signatures on the EEG signal and on the sleep wake-cycle architecture, providing mechanistic insights into its action and corroborating its traditional use to relieve sleeplessness, and favour relaxation.

## Results

To investigate the effects of *Tilia* extracts on sleep parameters, mice were fed an enriched diet for 3 weeks, followed by continuous EEG recordings over 5 days of treatment. The treatment did not alter the key electrophysiological features of wake, NREM or REM sleep (Fig. 1 A-B), allowing for accurate scoring of sleep-wake episodes (also indicated as behavioural states). Wakefulness was characterized by low-amplitude, high-frequency activity and irregular EEG signal; NREM sleep presented slow wave activity (SWA) characterized by high amplitude and slow frequency waves, while REM sleep exhibited a significant reduction of SWA, an increase in low-amplitude activity typically dominated by theta waves, and was easily differentiated from wake by the extremely reduced activity of the EMG traces as can be observed in the electrophysiological recordings and their relative spectrograms (Fig. 1A-C).

**Figure 1.**
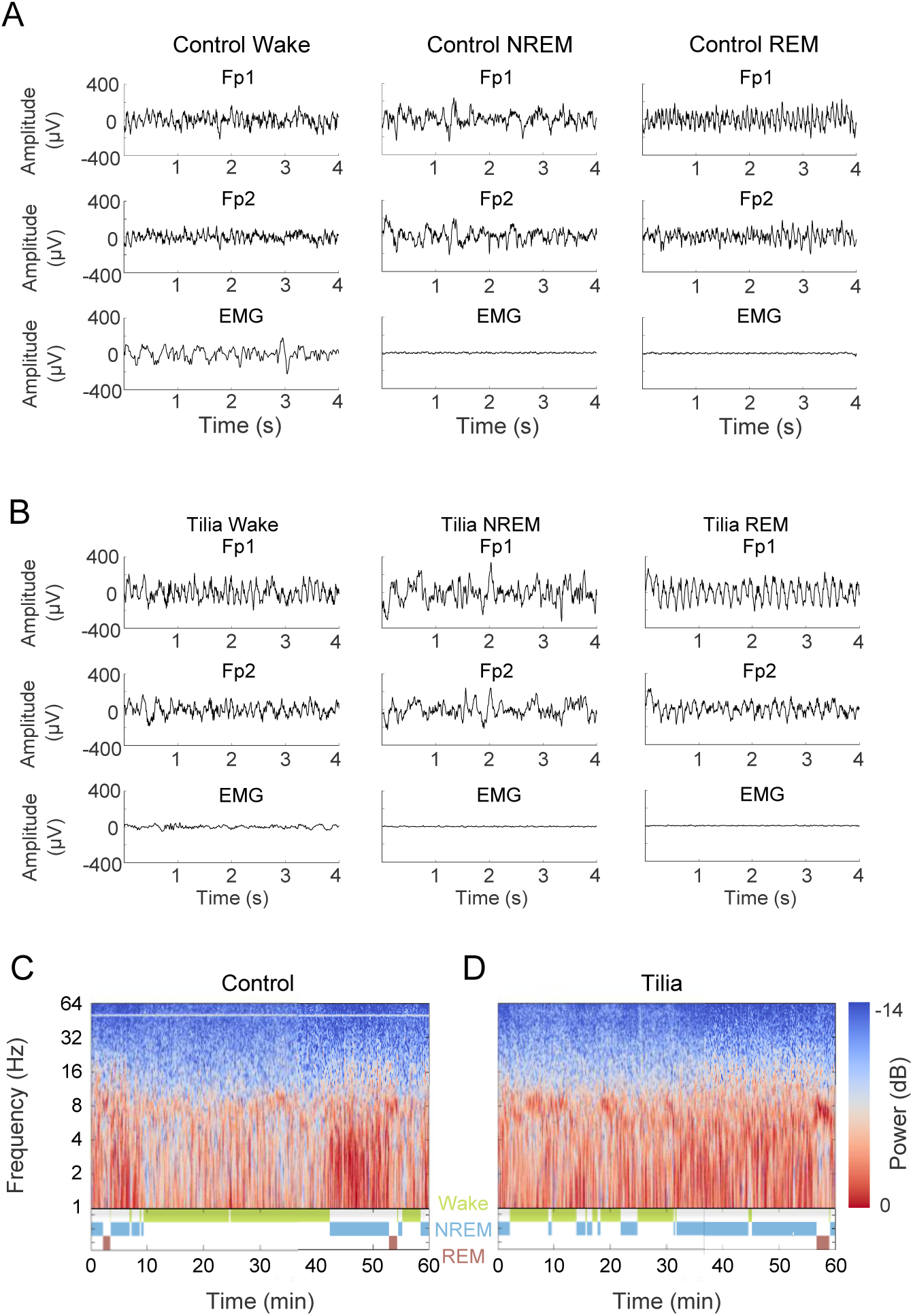
Wake, NREM, and REM sleep EEG signals in control and *Tilia-*treated mice. (A) Representative 4-second EEG epochs from a WT mouse during wake (left), NREM sleep (center), and REM sleep (right). Fp1 represents the signal recorded from the frontal lobe, Fp2 indicates the signal recorded from the parietal lobe, and EMG refers to the electromyogram signal. (B) Example 4-second EEG epochs from a mouse treated with *Tilia Tomentosa*. (C) Spectrogram showing changes in frontal EEG spectral power over one representative hour of recording in a WT mouse. (D) Spectrogram for a treated mouse. Each spectrogram is paired with its corresponding hypnogram, indicating the animal’s behavioural states over the same period. Changes in EEG power align with shifts in behavioural states.

We first analysed the effect of *Tilia* on the sleep-wake cycle architecture across the 24 hours. We observed that during the light phase (0-12 ZT), which reflects the resting period for nocturnal animals such as mice, the time course and percentage of wake and NREM sleep were similar in non-treated control and treated animals. When light was switched off, (at 12 ZT), controls exhibited a clear and considerable increase in wake, as expected in nocturnal animals. Concomitantly, they exhibited a substantial reduction in NREM sleep (Fig. 2A). Treated animals presented the same time course of these behavioural states, however, they had a smaller increase in wake and a less pronounced reduction in NREM sleep, during the same period, compared to controls (Fig. 2A and Fig. S1). In darkness (after ZT 19), controls showed a marked increase in wake and an equivalent reduction in NREM sleep, reflecting normal nocturnal activity. Noteworthy, *Tilia*-treated mice presented a reversal of this pattern, with a higher percentage of NREM sleep than wake (Fig. 2A and Fig. S1), during the same dark period (ZT 12-24) (Fig. 2 and Fig. S1). The time course and percentage of REM sleep appeared similar in the two groups of mice (Fig. 2A).

**Figure 2.**
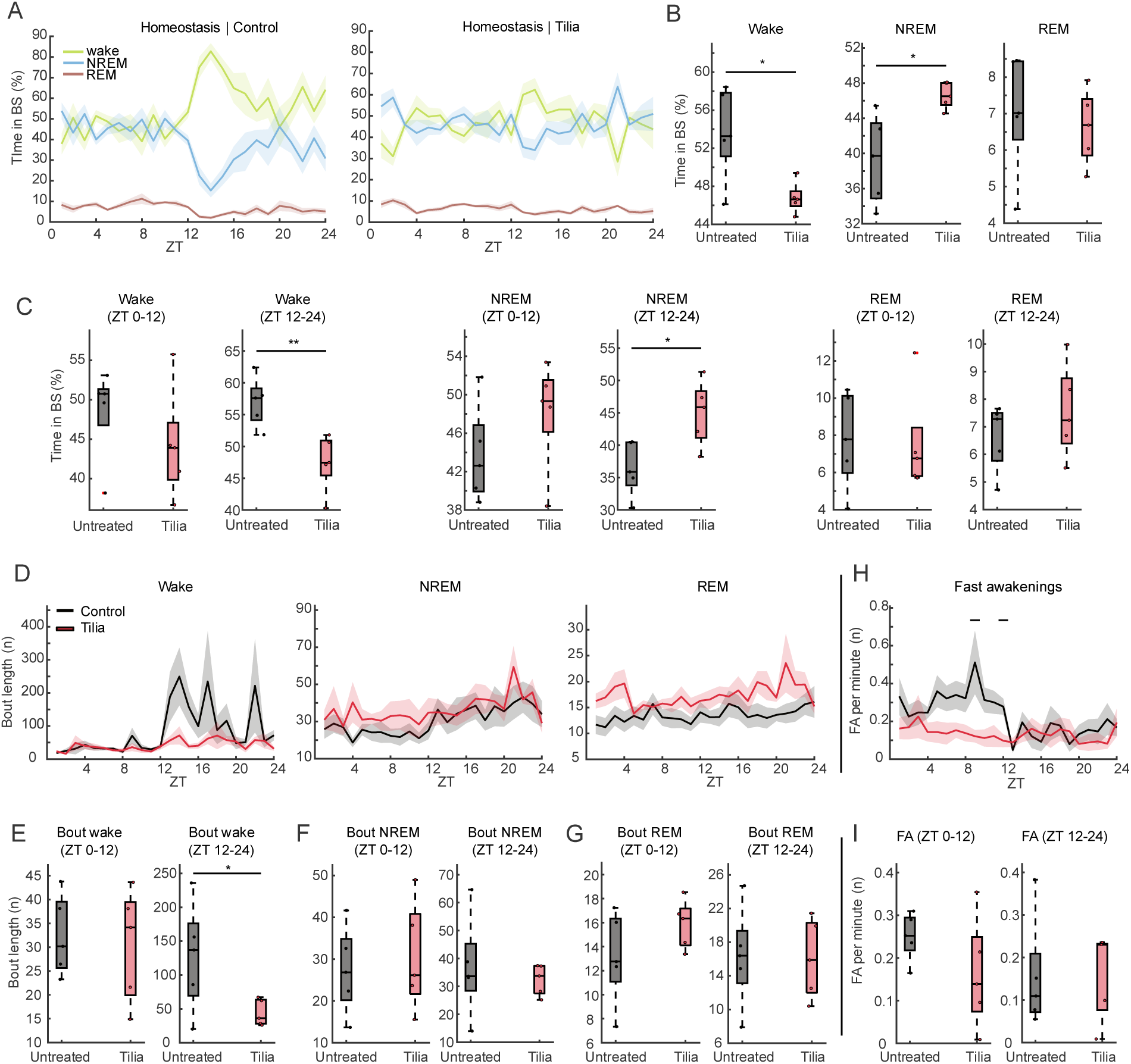
Sleep architecture in control and *Tilia*-treated mice. (A) Average percentage of time (lines) spent in each behavioural state across the 24 hours, calculated over five days of recording, for control (left) and *Tilia-*treated (right) mice. Shades indicate SEM. (B) Total percentage of time spent in wake (left), NREM sleep (center), and REM sleep (right) across 24 hours. Asterisks indicate statistical significance based on an unpaired t-test (*p < 0.05, **p < 0.01, ***p < 0.001). (C) Percentage of time spent in wake (left), NREM sleep (center), and REM sleep (right) during the light phase (ZT 0–12) and dark phase (ZT 12–24). (D) Bout length, indicated as number (n) of bouts, over the 24 hours for wake, NREM and REM sleep, respectively. Lines = mean. Shades = SEM (E–G) Bout length for wake (E), NREM sleep (F), and REM sleep (G) during the light and dark phases. (H) Number (n) of fast awakenings per minute across 24 hours. Solid black lines indicate statistical significance (p < 0.05). (I) Number (n) of fast awakenings per minute during the light and dark phases.

By quantifying the percentage of time spent in each behavioural state over the 24-hours, we found a significant reduction in wake and a corresponding increase in NREM sleep, in treated mice compared to controls. The amount of REM sleep was not significantly affected by the treatment (Fig. 2B). To assess whether *Tilia*’s effects varied across phases of the circadian cycle, we analysed the time course and the percentage of the different behavioural states during the light period (0-12 ZT) and dark period (12-24 ZT) separately (Fig. 2C). Although treated animals showed a trend towards reduced wakefulness and increased NREM sleep during the light period, these differences became statistically significant only during the dark period (12 -24 ZT segment). REM sleep appeared similar in both dark and light hours, in control and treated animals (Fig. 2C).

The quality of sleep depends not only on the total amount of sleep but also on its structure, i.e. if and how it is fragmented. To address this question, we measured the bout length for each behavioural state. A bout was defined as a sequence of at least four consecutive 4-second epochs in the same behavioural state. Therefore, bout length refers to the number of consecutive epochs within a given state. The analysis of this parameter, hour by hour, across the 24 hours, demonstrated that in controls, the duration of wake bout increased dramatically after lights went off (12 ZT), starting with the onset of the active phase. Another peak in wake bout length was observed in the central part of the dark phase, (around 21-22 ZT). Strikingly, this temporal evolution of wake bout duration was completely absent in mice fed a diet enriched with *Tilia* (Fig. 2D). In contrast, during the light phase – when mice are mostly resting – wake bouts were brief, with no appreciable differences between treated and control mice (Fig. 2 D). The bout length of NREM and REM sleep was similar in treated and control subjects (Fig. 2D). Statistical analysis conducted in the two temporal windows corresponding to the light (ZT 0-12) and dark (ZT 12-24) periods, respectively, confirmed that wake bout duration lacked the typical increase during the dark period in treated mice. These results are consistent with the increase in NREM and decrease in wake during darkness in treated mice, supporting a sedative-hypnotic effect of *Tilia*. Bout durations for NREM and REM sleep remained unaffected across both temporal windows (0-12 ZT and 12-24 ZT), in treated mice (Fig. 2D).

Another key feature that contributes to sleep fragmentation is the occurrence of brief awakenings. This term indicates episodes of wakefulness that last ≤ three epochs of 4 seconds each, occurring during NREM sleep bouts of at least one minute in duration. These episodes of awakenings are often accompanied by small body movements. Analysing the number of fast awakenings, hour by hour, across the 24 hours, we found that control mice showed a higher number of fast awakenings during the light phase (0-12 ZT), with a clear rise in number at the beginning of the resting phase. The number of fast awakenings remained high for several hours, with peaks around 8 and 10 ZT (Fig. 2E). Interestingly, *Tilia*-treated mice did not exhibit this rise in fast awakenings during the light phase (ZT 0-12), consistent with a sedative-hypnotic effect of *Tilia*. As expected, fast awakenings were almost absent during the dark phase, which corresponds to the most active period in mice (Fig. 2D). Therefore, sleep architecture in treated mice was characterized by shorter periods (bout) of wake during the dark phase (12 -24 ZT), and fewer fast awakenings during the light phase (0-12 ZT) with respect to controls, suggesting a circadian modulation of *Tilia*’s effect, which exerts its sedative-hypnotic effect reducing the length and number of awake periods, and favouring overall sleep increase and continuity.

To assess the impact of *Tilia* on the spectral characteristics of electrophysiological signals related to the sleep-wake states, we recorded the EEG from the frontal and the parietal lobe of each mouse. At first, we calculated the Power Spectral Densities (PSDs) in each channel for each behavioural state, across the 24 hours. We observed a significant decrease in gamma-band power (40-100 Hz) during wake in animals treated with *Tilia*, in both the frontal and parietal lobes (Fig. 3, Fig. S2). Interestingly, during REM sleep, we observed a clear increase in theta activity (around 8 Hz) in the frontal and in the parietal signal, in treated mice.

**Figure 3.**
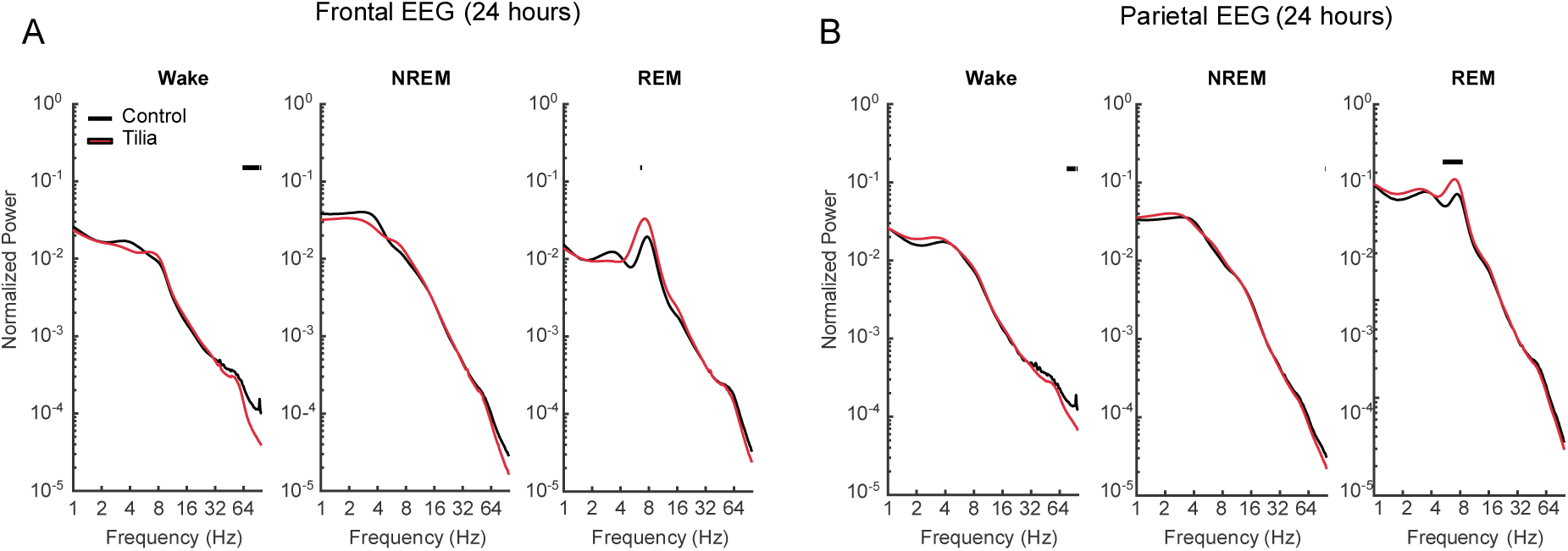
Spectral power in EEG signal over the 24 hours in control and *Tilia*-treated mice. (A) Average spectral EEG power in the frontal lobe for wake (left), NREM sleep (center), and REM sleep (right) across the 24 hours. (B) Same as (A), for the EEG signal recorded in the parietal lobe. Solid black lines indicate statistical significance (p < 0.05).

We then further dissected the effect of *Tilia*, performing spectral analysis of the EEG signal, separately examining the light (0-12 ZT) and dark (12-24 ZT) periods (Fig. 4). Interestingly, and coherently with the data reported above, most of the effects were observed during the dark period. In this time window, a significant decrease in gamma power was observed during wakefulness, in the signal recorded in both the parietal and frontal lobes, in treated mice. On the contrary, spectra of the REM sleep signal recorded in both areas presented a power increase in theta activity (around 8 Hz), in treated mice. Surprisingly, such theta increase during REM sleep was present also in the light period (0-12 ZT, Fig. 4).

**Figure 4.**
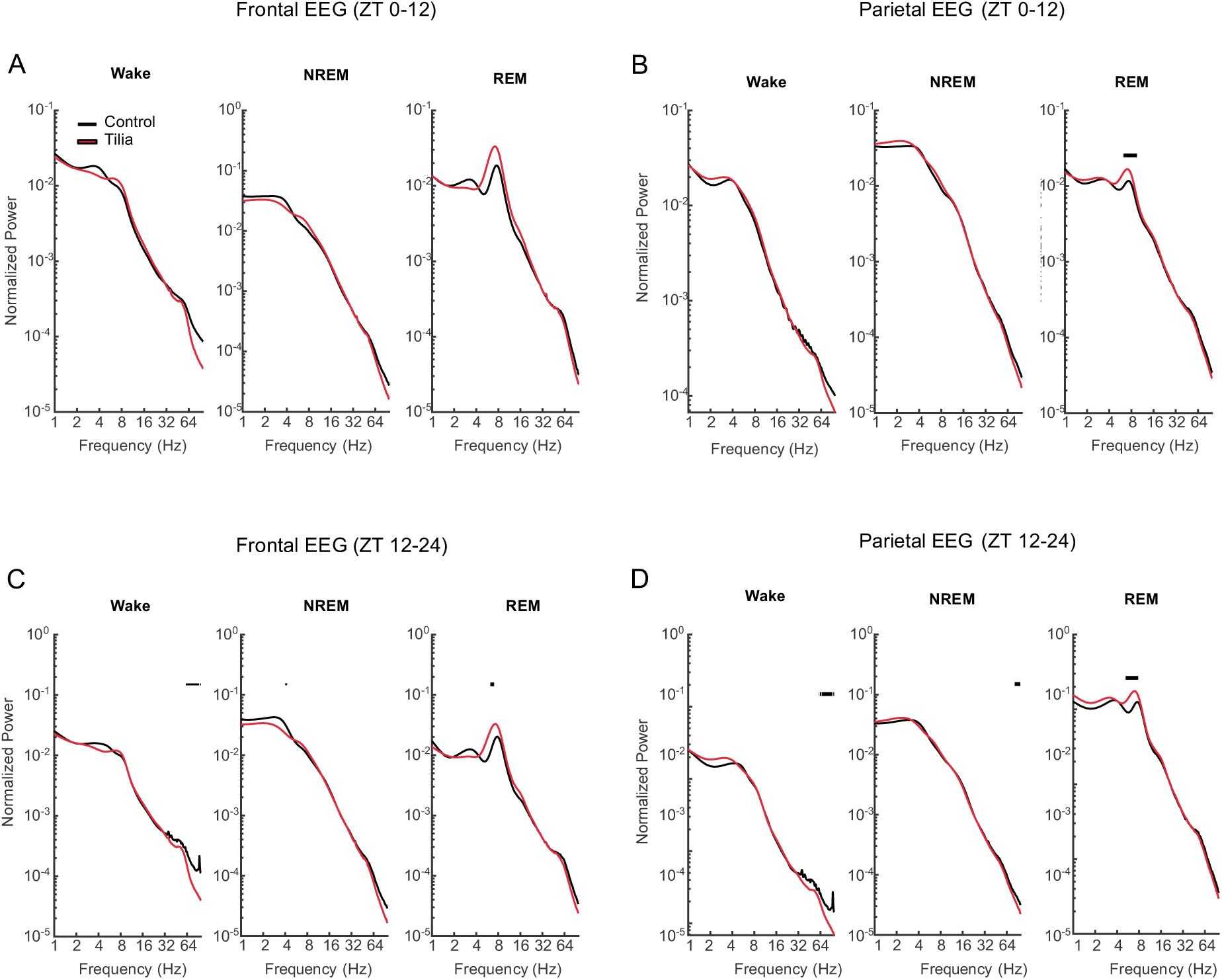
Spectral power in EEG signal during dark and light hours, in control and *Tilia*-treated mice. (A) Average spectral EEG power in the frontal lobe for wake (left), NREM sleep (center), and REM sleep (right) during the light phase. (B) Same as (A), for the EEG signal recorded in the parietal lobe. (C) Same as in A, for the EEG signal recorded during the dark phase. (D) same as in B, for the EEG signal recorded during the dark phase. Solid black lines indicate statistical significance (p < 0.05).

## Discussion

*Tilia* has been used for centuries in traditional medicine as a sedative remedy. More recently, several behavioural studies conducted in mice have provided compelling evidence of the tranquillizing and anxiolytic effects of Tilia extracts^6,12^. Treated mice appeared calmer, spent more time in quiet immobility, and were more prone to scout the environment around them and to engage in behaviours typically avoided by controls, such as entering and crossing the open arms of the T-maze. These data suggest reduced levels of anxiety with respect to controls and support the sedative action of *Tilia* on the central nervous system. Regarding the mechanism of action, electrophysiological studies performed in cultured hippocampal neurons demonstrated that *Tilia* operates via GABA receptors, strengthening the inhibitory tone, and reducing network excitability^10^. In particular, it was shown that flavonoids present in *Tilia* exert anxiolytic-sedative effects through GABA_A_ receptor activation, while Goutman et al. reported that flavonoids inhibit GABA receptors. This apparent contradiction can be explained by considering that flavonoids can exert anxiolytic effects via their antioxidant properties, which reduce the oxidative damage, without directly modulating GABA receptor currents.

Although the sedative and tranquilizing properties of *Tilia* are well-established, its effects on *in vivo* EEG signatures underlying the sleep-wake architecture and the circadian rhythm remained elusive. Here, we addressed these open questions by performing chronic EEG recordings in freely behaving mice, fed a diet enriched with *Tilia* extracts. We found that the treatment did not alter the typical features of the EEG signal, allowing us to promptly score the different behavioural states: wake, NREM and REM sleep. By analysing the time-course and the architecture of the wake-sleep cycle, we found that *Tilia* extracts induced a reduction in wake and a concomitant increase in NREM sleep, across the 24 hour. Interestingly, the increase in the amount of NREM sleep and the decrease in wake was particularly strong during the dark period, when mice are typically active. This phase-specific action suggests that *Tilia* promotes sleep mostly when the endogenous drive for wakefulness is high.

It has to be stated that, unlike humans, mice do not present a single, solid phase of sleep followed by a period of wakefulness. On the contrary, they alternate periods of sleep and wake along the 24 hours, being mostly active during the dark hours, and resting for most of the light hours. In this scenario, we found that *Tilia* significantly reduced the length of wake periods during the dark phase, i.e. during the period when mice are mostly active, but did not affect the length of wake bouts during the light phase, when animals spend most of the time resting. These results also point to a circadian-dependent action of Tilia. Importantly, the structure of NREM and REM sleep bouts remained unaffected, supporting the idea that *Tilia* promotes sleep by reducing arousal, rather than restructuring intrinsic sleep architecture.

The fact that *Tilia* extracts exert their strongest effects during the dark hours offers some particular advantages, specifically in circumstances in which there is the need of reducing movements and excitement in favour of relaxation, during the active phase. Human beings experience numerous situations in which a state of hyperactivity interferes with cognitive functions and performance. Therefore, interventions that can reduce arousal and promote relaxation may help improve attention and cognitive performance. The other striking advantage of *Tilia* extracts is its apparent lack of side effects, in contrast to conventional drugs with sedative and hypnotic effects, which are often associated with significant adverse effects. Benzodiazepines (BDZ), such as diazepam, are commonly prescribed to promote sedation and sleep. Several studies, conducted both in humans and animals, have shown that BDZs favour sleep by reducing sleep latency and increasing sleep continuity. However, they also tend to suppress REM sleep and alter network excitability, inducing distinct alterations of the brain’s physiological rhythmic activity^13^. At the level of the EEG spectra, BDZs lead to increases in fast-frequency activity, such as sleep spindles, and to reduction of the power of slow waves during NREM sleep. In addition, they decrease the incidence and the duration of OFF periods during NREM sleep^14^. Therefore, although BDZs facilitate sleep onset, they also alter the physiological structure of the network activity underpinning NREM and REM sleep^14–16^, suggesting that BDZs induce sleep with different characteristics with respect to the physiological one. These alterations could also contribute to the cognitive impairments often reported by individuals with insomnia who rely on BDZs to manage sleep disturbances. BDZs, finally, are also highly addictive, and patients using them on a regular basis often develop tolerance and substantial withdrawal symptoms, which can lead to their misuse. Because of these limitations, their use in the management of chronic sleep disturbances is generally discouraged^17–19^.

In contrast to BDZs, *Tilia* offers a valid alternative as a tranquillizer and hypnotic agent, as it modulates sleep patterns and neural activity without inducing structural alterations or abnormal EEG rhythms. In particular, the observed decrease in gamma activity during wakefulness and the increase in theta power during REM sleep are consistent with a sedated neural state. This stands in contrast to EEG spectral alterations typically associated with BDZs use, which often involves reductions in slow-wave activity and disruptions of natural sleep rhythms. Indeed, spectral analysis of the EEG recorded in treated mice presented a significant reduction in power in the gamma band during wakefulness, likely reflecting the reduced activity observed at behavioural level. Likewise, EEG spectra of REM sleep exhibit a clear increase in the theta range, suggesting a more coordinated activity of the network during this stage of sleep. Noteworthy, the effect on the EEG spectra were stronger during the dark phase, suggesting a circadian-specific action of Tilia.

Altogether, these data indicate that *Tilia* promotes relaxation and sleep without disrupting the physiological cortical network dynamics. In particular, slow-wave activity during NREM sleep remained stable, further indicating that *Tilia* does not impair the restorative features of deep sleep, unlike many conventional sedatives - such as benzodiazepines - have been shown to reduce slow-wave power (0-4 Hz) during NREM sleep^20^. Thus, *Tilia* appears to maintain the restorative deep-sleep component that is often compromised by standard hypnotics. Furthermore, up to now, no evidence has been reported regarding addiction associated with *Tilia* extracts.

Beyond GABAergic modulation, future investigations may also consider the potential role of serotonergic pathways in mediating *Tilia*’s effects. Several flavonoids found in *Tilia*, including quercetin, rutin, and kaempferol, are known to interact with serotonin receptors such as 5-HT1A and 5-HT2A/2C, which are implicated in the regulation of mood, arousal, and sleep. Exploring these alternative mechanisms could provide a more comprehensive understanding of *Tilia*’s neuropharmacological profile.

Altogether, our findings indicate that *Tilia* acts as a sedative agent that – overall – preserves the physiological architecture of sleep and EEG spectral profiles. These results support the potential use of *Tilia* for managing conditions such as restlessness, hyperactivity and insomnia.

## Methods

### Animals

*Animals* (control, n=5; treatment, n=5) were housed in filtered cages in a temperature-controlled room with a 12/12-h dark/light cycle (light on at 6 AM). All procedures were conformed to the EU Directive 2010/63/EU for animal experiments and to the ARRIVE guidelines. Experimental protocols were approved by the Italian Ministry of Health. Experiments were performed on female and male wild-type (C57BL/6J, The Jackson Laboratory, Stock No: 000664), at the age of about 8 weeks. All analyses were performed blind to the experimental condition.

### Dietary regimen

Mice were divided in 2 groups: 1) one group had access to 4 g/die of regular food; 2) the other group were fed 4 g/die food enriched in *Tilia’s* extracts. Both groups had access to water *ad libitum.* Standard food for mice was enriched with fluid extracts of *Tilia:* 3 g of standard food contained 2.34 ml of fluid extracts of *Tilia*.

### Surgical procedure

Animals were anesthetized with a mixture of Zoletil100 (a combination of Zolazepam and Tiletamin, 1:1, 10 mg/kg; Virbac, Carros, Cedex, France) and Rompun (Xylazine 2%, 0.06 mL/kg; Bio98, Milan, Italy) and placed into a custom stereotaxic frame. Two small holes (∼1.2 mm diameter) were made in the skull to allow the insertion of two iron-plated, round-tipped miniature screws that served as EEG electrodes. Electrodes were placed over the parietal lobe (3 mm lateral to midline, 2 mm posterior to bregma) and the frontal lobe (1.5 mm lateral to midline, 1.5 mm anterior to bregma). A third screw, used as a reference electrode, was fixed on the skull above the cerebellum (2 mm posterior to lambda, on the midline). Two small stainless steel wires (Advent, Stock No: FE635011) were inserted into the neck muscles and served to record the electromyogram (EMG). Electrodes were soldered to stainless steel wires and secured to the skull with dental cement (Paladur, AgnTho’s, Lidingö, Sweden). Mice were immediately treated with post-operative analgesia (Tramadol 10 mg/kg, Formevet, Milan, Italy). Upon waking up from the anaesthesia, animals were placed in individual cages for 5 days to allow complete recovery before initiation of EEG recordings.

### EEG acquisition and analysis

The electroencephalographic signal was recorded using the Micromed Brain Quick LTM Holter EEG system (Micromed, Mogliano Veneto, Italy). Data were acquired sampled at 256 Hz. Mice were free to move in a plexiglass arena (35 cm × 35 cm × 40 cm) placed within a Faraday cage. Before the beginning of the recording session, mice were habituated to the apparatus for about 30 min. The three behavioural states - wake, NREM and REM sleep - were determined for 4-s epochs by visual inspection of the EEG and EMG signals using the Sirenia Sleep software (Pinnacle Technology). Epochs containing artifacts in any one of the recorded channels were automatically excluded from the subsequent analyses. After the manual scoring, the recorded signal altogether with the information about the behavioural states were imported on MATLAB (MathWorks Inc.) and further analysed. We considered ‘bouts’ as periods of at least three consequent 4-s epochs that mice spent in a single behavioural state. Power spectral density (PSD) of EEG was computed over each epoch time interval with Welch’s method, choosing a Hann window with a length of 0.5 s and with 0.25 s overlap between windows. Normalized power was calculated dividing the power in each behavioural state over an hour of recording by the power of the whole recording, irrespective of the behavioural state.

### Statistics

Statistical analyses were conducted using the built-in Matlab function *ranksum* to perform Wilcoxon tests. Differences in power spectra were assessed using independent t-tests at each discrete frequency value, applying a significance threshold of p < 0.05. No correction for multiple comparisons was applied, in line with common practice, as such corrections are typically overly conservative when comparing spectra across many frequency bins. All tests were two-sided. Sample sizes were chosen to reflect standard practices in exploratory *in vivo* rodent electrophysiology studies; no formal power calculations were performed.

## Acknowledgments

We thank Marco Brondi for insightful suggestions, and Beatrice Perissutti, for preparing the food. Work by CL supported and funded by: #NEXTGENERATIONEU (NGEU); the Ministry of University and Research (MUR); the National Recovery and Resilience Plan (NRRP); project MNESYS (PE0000006) – A Multiscale integrated approach to the study of the nervous system in health and disease (DN. 1553 11.10.2022). This work was also supported by the Veneto Regional grant POR – FESR - Action 1.1.4., 3S_4H.

## Authors Contributions

CL design the research; ADS and MC performed the experiments; ADS performed data analysis with help from MC; NR contribute to optimization of the hardware for the EEG recordings (Micromed Brain Quick LTM Holter EEG system (Micromed, Mogliano Veneto, Italy)); CL supervised the experiments and data analysis. CL and ADS wrote the paper that was read and approved by all the authors.

## Competing interests

The authors declare no competing interests.

**Correspondence** and requests for materials should be addressed to Claudia Lodovichi

## Figure legends

**Figure S1.**
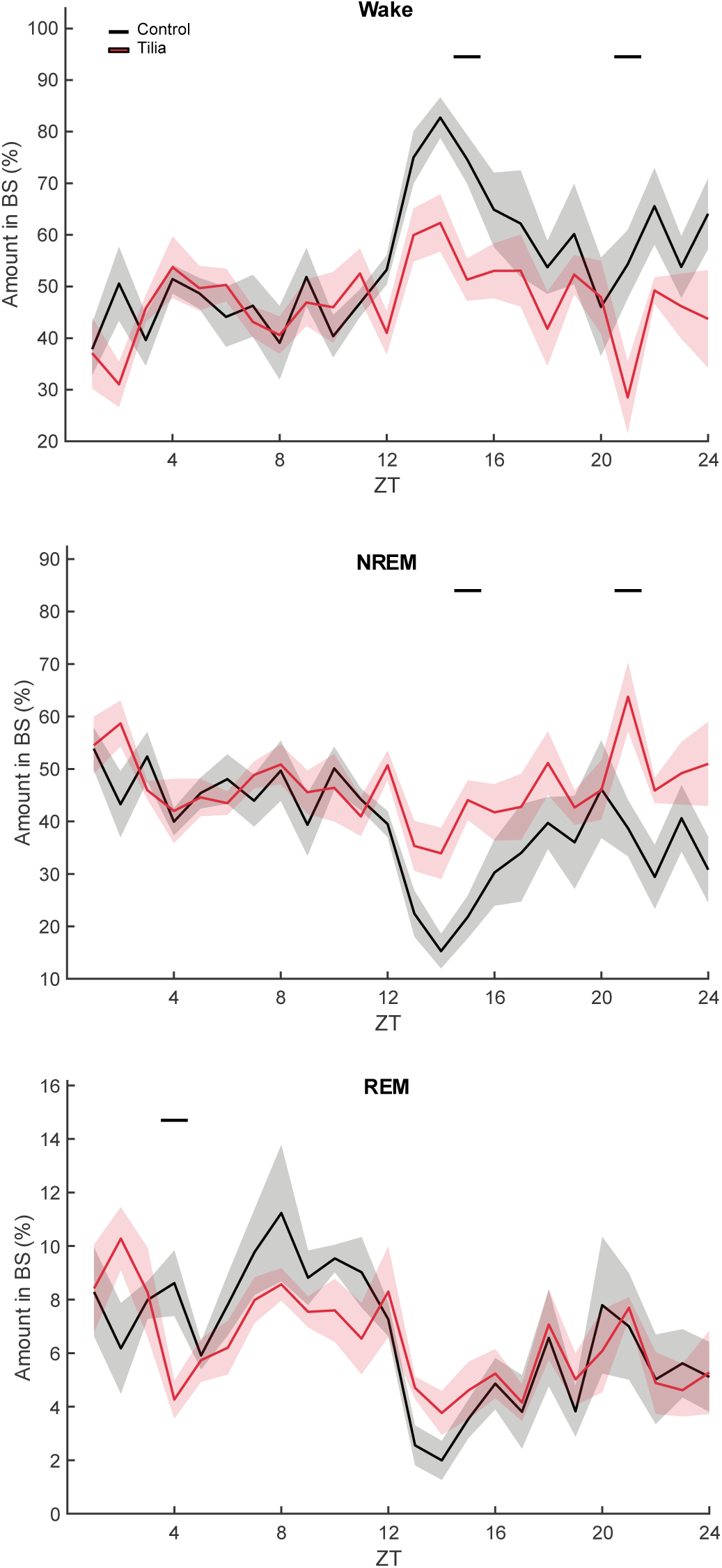
Wake, NREM and REM sleep in control and *Tilia*-treated mice. (A) Average percentage of time of wake (top), NREM sleep (center) and REM sleep (bottom) across the 24 hours, calculated over 5 days of recording. Lines = mean. Shades = SEM. Solid black lines, on top of graphs, indicate statistical significance (p < 0.05).

**Figure S2.**
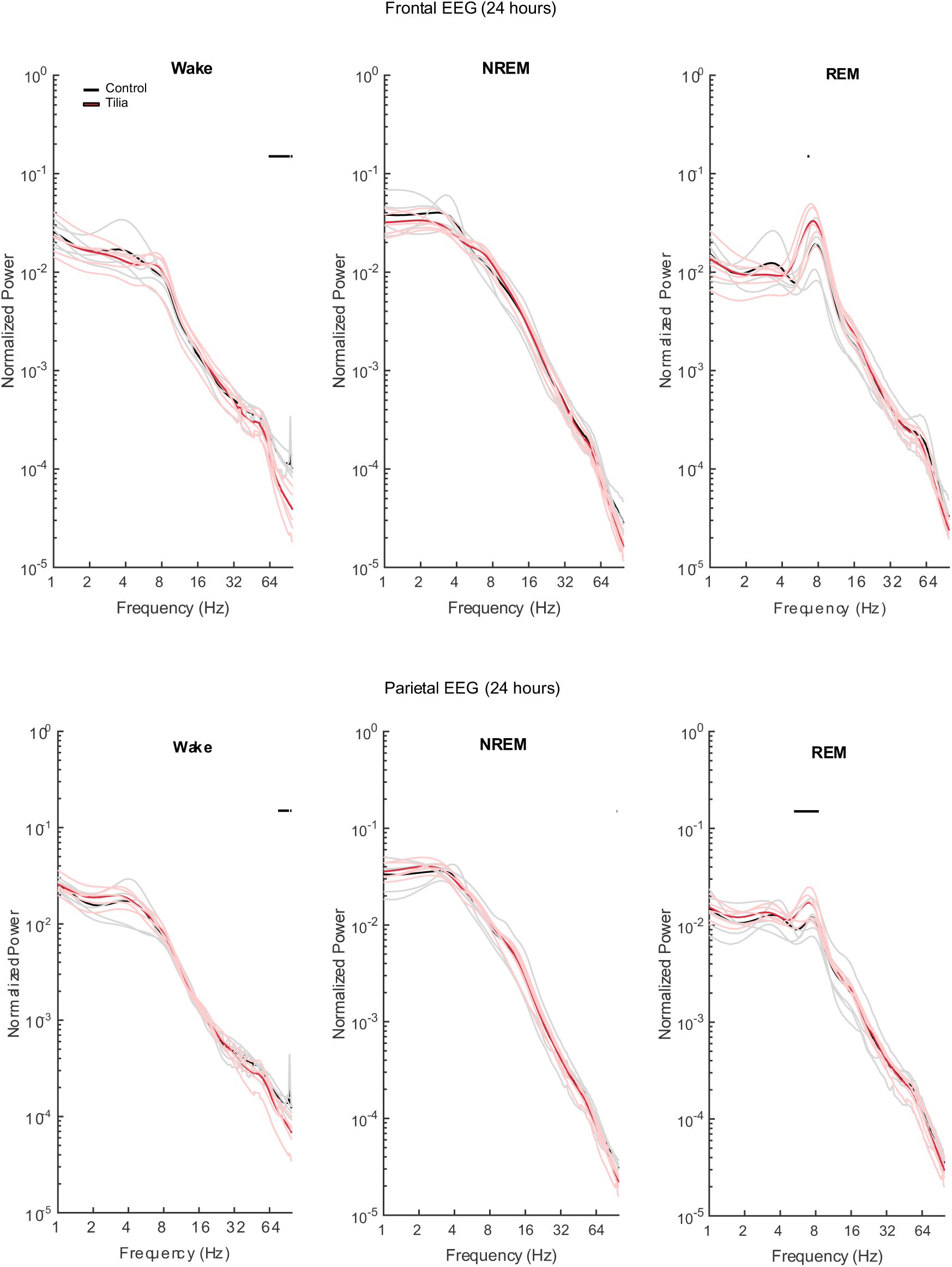
Spectral power in EEG signal over the 24 hours in control and *Tilia*-treated mice. (Top row) Average spectral EEG power in the frontal lobe for wake (left), NREM sleep (center), and REM sleep (right) across the 24 hours. (Bottom row) Same as (A), for the EEG signal recorded in the parietal lobe. Spectral power is normalized by dividing the power for each behavioral state by the total spectral power of the entire recording, regardless of behavioral state. Solid black lines indicate statistical significance (p < 0.05). Thin lines indicate single mice.

